# A multi-faceted analysis of synapses reveals the role of neuroligin-1 cleavage in presynaptic vesicle accumulation in the lateral amygdala

**DOI:** 10.1101/2023.11.07.566075

**Authors:** Connon I. Thomas, Jordan R. Anderson, Arman Alexis, Debbie Guerrero-Given, Abigail Chavez, Micaiah C. McNabb, Bengi Unal, Michael D. Ehlers, McLean M. Bolton, Naomi Kamasawa

## Abstract

Neuroligin-1 (NLGN1) is a cell adhesion molecule found at excitatory glutamatergic synapses in the brain which regulates synaptic function and maturation. Extracellular cleavage of NLGN1 by proteases has been shown to control vesicle release in cultured neurons, but nothing is known about the underlying changes to synapse structure that accompany this, or how synapse function is affected in brain tissue. We found that prevention of NLGN1 cleavage through mutation to the extracellular stalk domain increases synaptic vesicle docking and miniature excitatory post-synaptic current frequency at synapses of the lateral amygdala. Using a novel volume electron microscopy (vEM) analysis pipeline based on deep learning extraction of thousands of synapses and vesicles clouds and subsequent spatial analyses, we found that the total pool of synaptic vesicles shifts closer to the synapse in mutants. Furthermore, we observed an increased frequency of incomplete synapses that lack vesicle accumulation, pointing towards disruption of synaptic pruning and accumulation of putatively non-functioning synapses. Our study provides evidence of a structural and functional role of NLGN1 cleavage in native brain tissue, and establishes a foundation for vEM analysis of synapse-vesicle spatial relationships in other animal models of dysfunction and disease.

## Introduction

Neurons in the brain communicate at specialized junctions called synapses. Cell-adhesion molecules (CAMs) structurally link pre- and postsynaptic membranes, and often have other functions through interactions with synaptic machinery. Neuroligin-1 (NLGN1) is one such adhesion molecule; located at the postsynaptic membrane of excitatory synapses, it is critical across development in shaping glutamatergic synapse function through structural modification of the synapse^1–3^. As NLGN1 mutations are associated with Autism Spectrum Disorder (ASD) and Alzheimer’s Disease, developing a holistic understanding of its modes of action is of critical importance^3–5^.

Both the structure of NLGN1 and the interactions it has with other synaptic molecules are well studied. It is composed of several distinct structural domains: an N-terminal acetylcholinesterase domain (AChE), a juxtamembrane stalk, a single transmembrane-spanning region, and a C-terminal PDZ binding domain^6^. Through these domains, NLGN1 couples presynaptic neurexins to postsynaptic scaffolding molecules, allowing for NMDA and AMPA receptor recruitment and alignment of pre- and postsynaptic machinery^7–17^. NLGN1 is not static at the synapse but is cleaved by proteases, including matrix metalloprotease-9 (MMP9) and ADAM10, at the juxtamembrane stalk region. This cleavage releases the N-terminal ectodomain and destabilizes its presynaptic partner, neurexin-1β^18,19^. N-terminal cleavage has been demonstrated to occur at individual spines, is triggered by synaptic activity, and is dependent on NMDA receptor and Ca^2+^/calmodulin-dependent protein kinase^18^. In cultured hippocampal neurons, the acute effect of NLGN1 ectodomain cleavage is a rapid decrease in release probability with no change in postsynaptic responsiveness^18^, whereas, chronic overexpression of the n-terminal ectodomain promotes spine formation^19^.

Understanding the role of NLGN1 cleavage in vivo and in native brain tissue is of increasing interest given the prevalence of activity dependent cleavage of CAMs by proteases and its implications in disease. For example, MMP9 induces spine enlargement and is required for hippocampal LTP^20–23^, but is also a key player in pathological remodeling in disease processes such as in epileptogenesis, stroke, cancer, and inflammation^24,25^. Despite evidence that NLGN1 ectodomain cleavage by proteases occurs, the role of this cleavage and the resulting cleavage fragments in regulating basal synaptic structure and function in brain tissue has yet to be characterized.

While electrophysiology can be used to probe the functional properties of synapses, the gold-standard for ultrastructural measurement is electron microscopy (EM). Thin-section transmission EM (TEM) can be used to quantify changes to synaptic morphology at high resolution; however, the three-dimensional shape and size of synapse components, including presynaptic vesicle pools, are most easily obtained using volume EM (vEM) techniques such as serial block-face scanning EM (SBF-SEM). A limitation of performing vEM is the “analysis bottleneck”—following acquisition of massive image datasets, segmentation and quantification can take months to years to complete^26,27^. However, in recent years, the development of deep learning (DL) methods has increased the throughput of the segmentation pipeline^28–31^. Despite the importance of vesicle accumulation and synapse morphology in determining synaptic function, little has been done to develop methods for characterizing synapse and vesicle structure in EM volumes, particularly the spatial relationship between the two.

In this study, we characterized the function and structure of synapses in amygdala tissue of mice bearing a mutation in a portion of the NLGN1 juxtamembrane stalk called the SD3 region that prevents its cleavage. We found that miniature excitatory postsynaptic current (mEPSC) frequency is enhanced, more presynaptic vesicles are docked per synapse, and vesicles are overall slightly closer to the synapse. We developed a novel vEM analysis pipeline based on deep learning extraction of synapses and vesicles, pairing of synapse and synaptic vesicle clouds (SVC), measurement of their alignment to one another, and automated calculation of distances between them. Using this pipeline, we observed that mutants had a greater percentage of synapses with an SVC in contact with the synapse. Furthermore, mutants showed an increased frequency of “incomplete” synapses which lack vesicle accumulation, pointing towards disruption of synaptic pruning and accumulation of putative synaptic remnants. Our study provides evidence of structural and functional disruption of synapses caused by loss of NLGN1 cleavage, and establishes a foundation for vEM analysis of synapse-vesicle spatial relationships in other mutant models.

## Results

### Utilization of a multi-faceted approach to characterizing synapse structure and function in the lateral amygdala

We chose to focus our study on the lateral amygdala due to the requirement of NLGN1 for normal fear learning, NMDA receptor expression and synaptic plasticity in the adult lateral amygdala, and importance of the amygdala in regulating social and emotional behavior in ASD^32,33^. To provide a more comprehensive understanding of how NLGN1 cleavage affects synapse structure and function, we combined electrophysiology, TEM, and SBF-SEM techniques. Extraction of features from vEM data requires segmentation on a large scale, involving massive amounts of human labor. We therefore increased our throughput using DL networks to efficiently extract thousands of synaptic features from sets of volumetric image data. We then developed a toolset and pipeline designed to assess changes in the aggregation of synaptic vesicles at the synapse and survey hundreds to thousands of features in the tissue.

### Deep learning based 3D segmentation of synapses and vesicle clouds

We trained two sets of models to segment our SBF-SEM data—one for synapses and one for vesicle clouds—using integrated DL architectures available within the ORS Dragonfly software. To prepare training data sets, synapses were manually demarcated as the presynaptic membrane, rigid synaptic cleft, and postsynaptic membrane. Networks that use slice-to-slice context have been shown to have higher accuracy^34^, therefore we chose to train 2.5D and 3D architectures to detect features using several sections of context, as synapses and vesicle clouds can be tracked through approximately 3-10+ sections in our datasets (Figure 1A-C, details are described in Methods).

**Figure 1:**
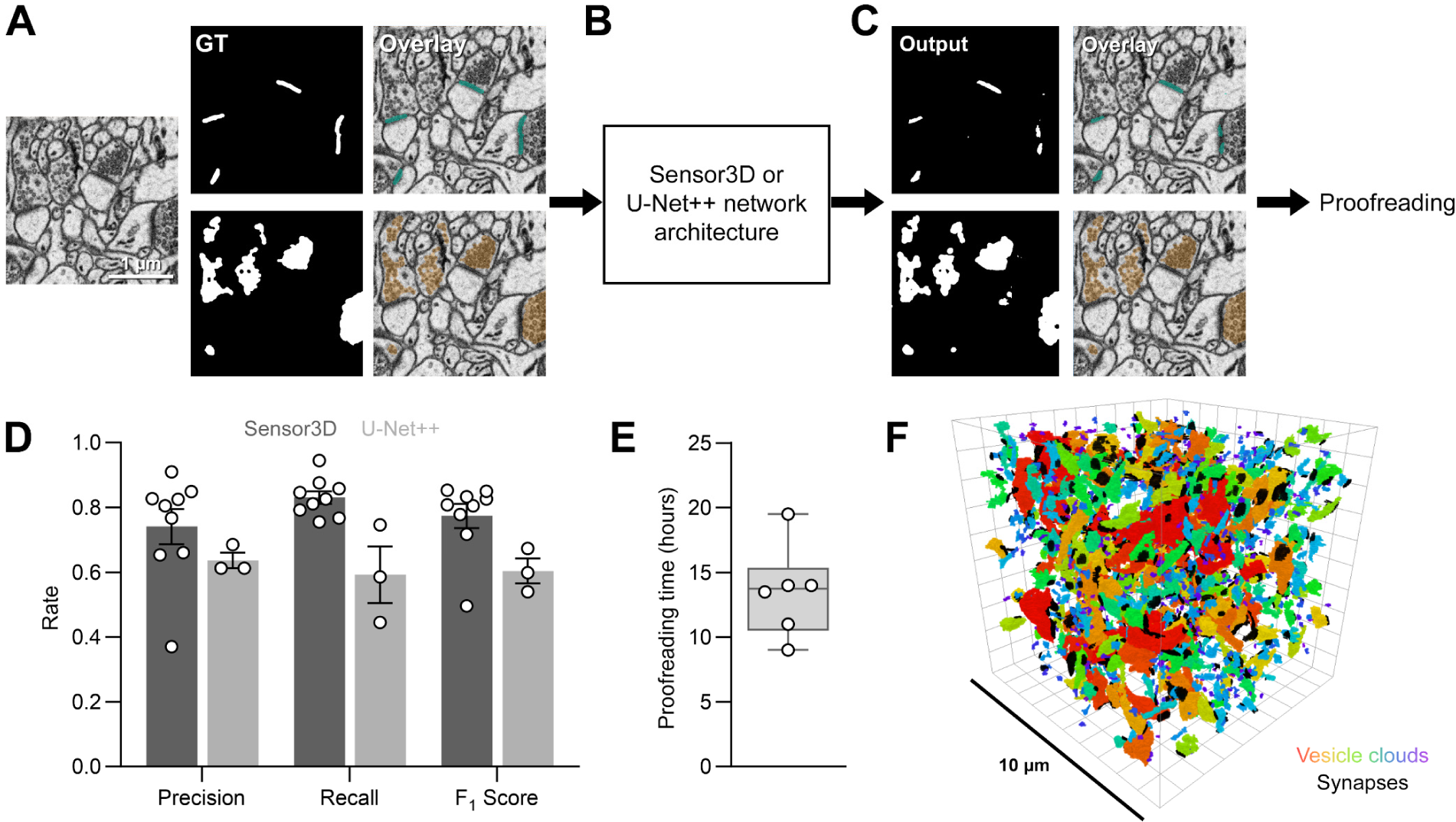
Automated segmentation workflow and network performance. **A.** Example SEM image, binary synapse (cyan) and vesicle (orange) segmentation ground truth (GT) labels, and overlay. **B.** Ground truth segmentations were fed into one of two deep learning network architectures. **C.** Trained network output binary and overlay. The network used for this segmentation was not trained on the ground truth data shown in A. **D.** Network segmentation performance. Synapse and vesicle networks are pooled (n = 12 networks). **E.** Time required for proofreading following automated segmentation of 10,000 µm^3^ volumes. **F.** 3D rendering of synapses and vesicles from a 10,000 µm^3^ volume. Vesicle clouds are colored by size (mutant amygdala, n = 552 synapses, 1268 vesicle clouds). Error bars represent mean ± S.E.M.

We evaluated the performance of our models using standard metrics including the precision rate (the ratio of correctly predicted positive observations to the total predicted positives), recall rate (the ratio of correctly predicted positive observations to all the actual positives), and F1 score (the weighted average of precision and recall) for each network we trained (**Figure 1D**). We calculated a median F1 score of 0.599 across U-Net++ models and 0.808 across Sensor3D models, suggesting that our Sensor3D models performed considerably better. We observed that the poorest performing networks were generally the first that we trained, and the best performing networks were retrained several times on new data. To maximize the quality of our segmented dataset, outputs were proofread by an experienced annotator as described in the methods before generating final renderings (**Figure 1E, F**). Altogether, we extracted 2855 synapses and 4719 vesicle clouds from all six animals.

### Toolset development for 3D measures of vesicle geometry in vEM data

In order to investigate the relationship between vesicles and synapses in a volume with a considerable sample size (hundreds to thousands of features), as well as measure potential changes in vesicle cloud geometry and spatial organization relative to the synapse, we needed to develop an efficient analysis pipeline. This pipeline involves pairing synapses with vesicle clouds based on their biological relevance, followed by automated quantification of vesicle cloud offset relative to the synapse. We created openly available, python-based plugins for Dragonfly (see methods) that extract batches of synapse and vesicle cloud meshes, pair meshes based on nearest-neighbor mesh distance, and quantifies synapse-vesicle cloud arrangement (**Figure 2A**).

**Figure 2:**
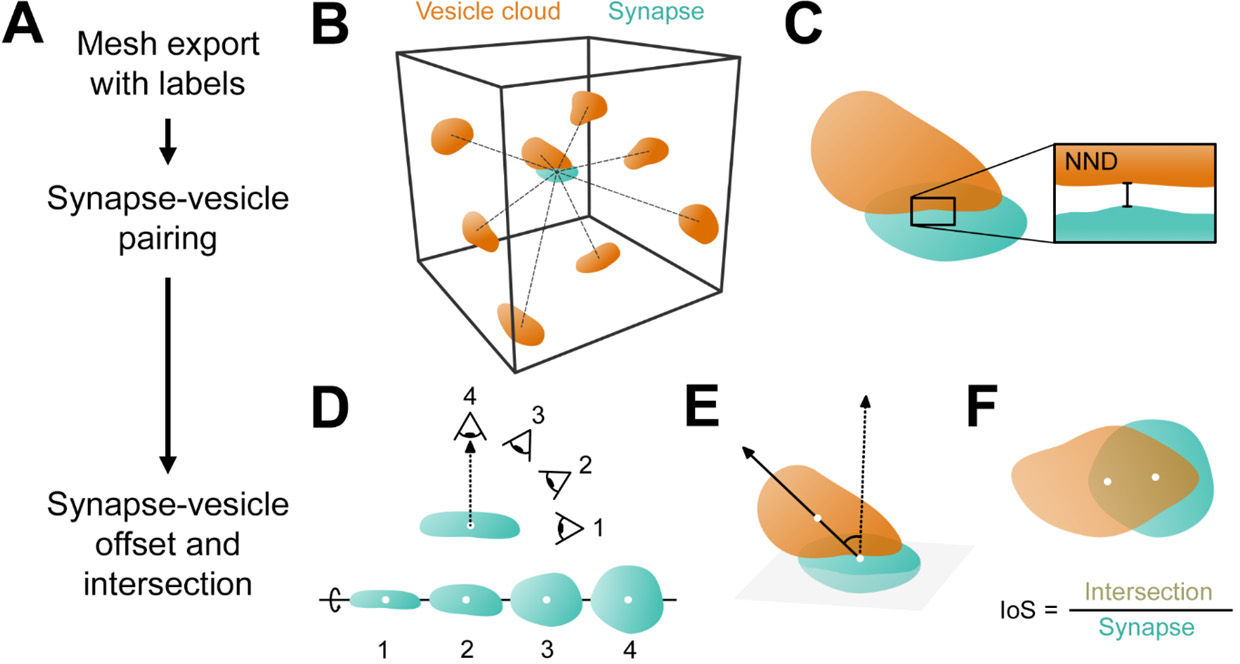
3D synaptic analysis pipeline. **A.** Pipeline overview. Following completion of proofreading, meshes were exported from Dragonfly in order of size. Synapses were then paired to vesicle clouds, and cloud offset and intersection were measured. **B.** For a given synapse mesh, the center of mass (CoM) distance was calculated to all vesicle cloud meshes within a 2 µm search window. These clouds were considered candidate pairs. **C.** The synapse mesh was paired to the candidate vesicle cloud with the shortest nearest-neighbor distance (NND). **D.** The maximum projection (i.e. #4) for a synapse was found using a virtual camera that rotates around the synapse CoM (white dot). The line connecting the virtual camera and the synapse CoM (dotted line) represents the vector normal to the synaptic plane, and was used for offset angle and intersection measurements. **E.** Offset angle measurement between the synapse-normal vector and vector that connects the vesicle cloud CoM to the synapse CoM. **F.** Intersection measurement, represented as the intersection of the SVC projection and synapse maximum projections, divided by the synapse maximum projection.

We sought to pair each synapse with the vesicle cloud in its presynaptic compartment. However, without extraction of the presynaptic cell membrane, compartmental information for each vesicle cloud was lost. Since the majority of synapses have docked vesicles or vesicles within tens of nanometers from the active zone^35^, we chose to use the distance between the synapse and vesicle cloud as a method of pairing vesicle clouds to the synapse where release presumably occurs. For each synapse mesh, we identified all vesicle clouds with centers of mass within a 2 µm cube to increase computational speed (**Figure 2B**). We then calculated the nearest neighbor distance (NND) based on mesh vertex distances, and assigned the vesicle cloud with the shortest mesh-mesh distance as its pair (**Figure 2C**).

The final step of our pipeline was developed to capture the spatial offset and misalignment of the vesicle cloud relative to the synapse. Using the PyVista library and a virtual camera system, we collected projections of each synapse from different positions arranged radially around the center of mass (CoM) of the synapse (**Figure 2D**). The viewpoint that yielded a 2D projection of a synapse with the greatest surface area was considered the viewpoint that exists on a vector that is roughly normal to the synaptic plane. We then calculated vesicle offset as the angle between the vector normal to the synaptic plane and the vector that passes through the synapse CoM and vesicle cloud CoM (**Figure 2E**). For the alignment measurement, we used the virtual camera system to capture the area of overlap between projections of a synapse (viewed from the normal vector) and its associated vesicle cloud. We calculated the intersection of the vesicle and synapse projections, and divided this by the synapse projection area. We termed this IoS, or intersection over synapse (**Figure 2F**).

### Prevention of NLGN1 cleavage increases vesicle release probability at excitatory synapses in the lateral amygdala

Previous investigation in cultured hippocampal neurons has shown that when NLGN1 cleavage is blocked by expressing the cleavage-resistant mutant NLGN1-ΔSD3, vesicle release probability increases^18^. It is unclear whether this effect applies in vivo, in other neuron classes, or different brain areas. We therefore probed whether the same mutation introduced in knock-in mice affects mEPSC frequency and amplitude in the LA and measured vesicle proximity to the synapse using thin-section TEM and SBF-SEM.

We found no difference in the mean mEPSC amplitude between WT and NLGN1-ΔSD3 mutant mice (mean: WT = 11.28 ± 0.59 pA, n = 19 cells; Mutant = 12.46 ± 0.90 pA, n = 21 cells, p = 0.2796, t-test), but found that mutants had significantly higher mEPSC frequency (mean: WT = 0.97 ± 0.15 Hz, n = 19 cells; Mutant = 1.86 ± 0.22 Hz, n = 21 cells, p = 0.0024, t-test; **Figure 3A-C**). This suggests while the mean size of events did not change, synaptic vesicle release probability may be increased in NLGN1-ΔSD3 mutant mice. When looking at the pooled amplitude cumulative distribution, we observed a significant shift towards larger amplitudes, particularly in the upper half of samples (median: WT = 10.99 pA, IQR = 5.42 pA, n = 19 cells; Mutant = 11.73 pA, IQR = 7.31 pA, n = 21 cells, p < 0.0001, MW test; **Figure 3D).** We also observed a significant decrease in the cumulative distribution of mEPSC interspike intervals in mutants (median: WT = 0.82 Hz, IQR = 1.72 Hz, n = 19 cells; Mutant = 0.41 Hz, IQR = 0.68 Hz, n = 21 cells, p < 0.0001, MW test; **Figure 3E**).

**Figure 3:**
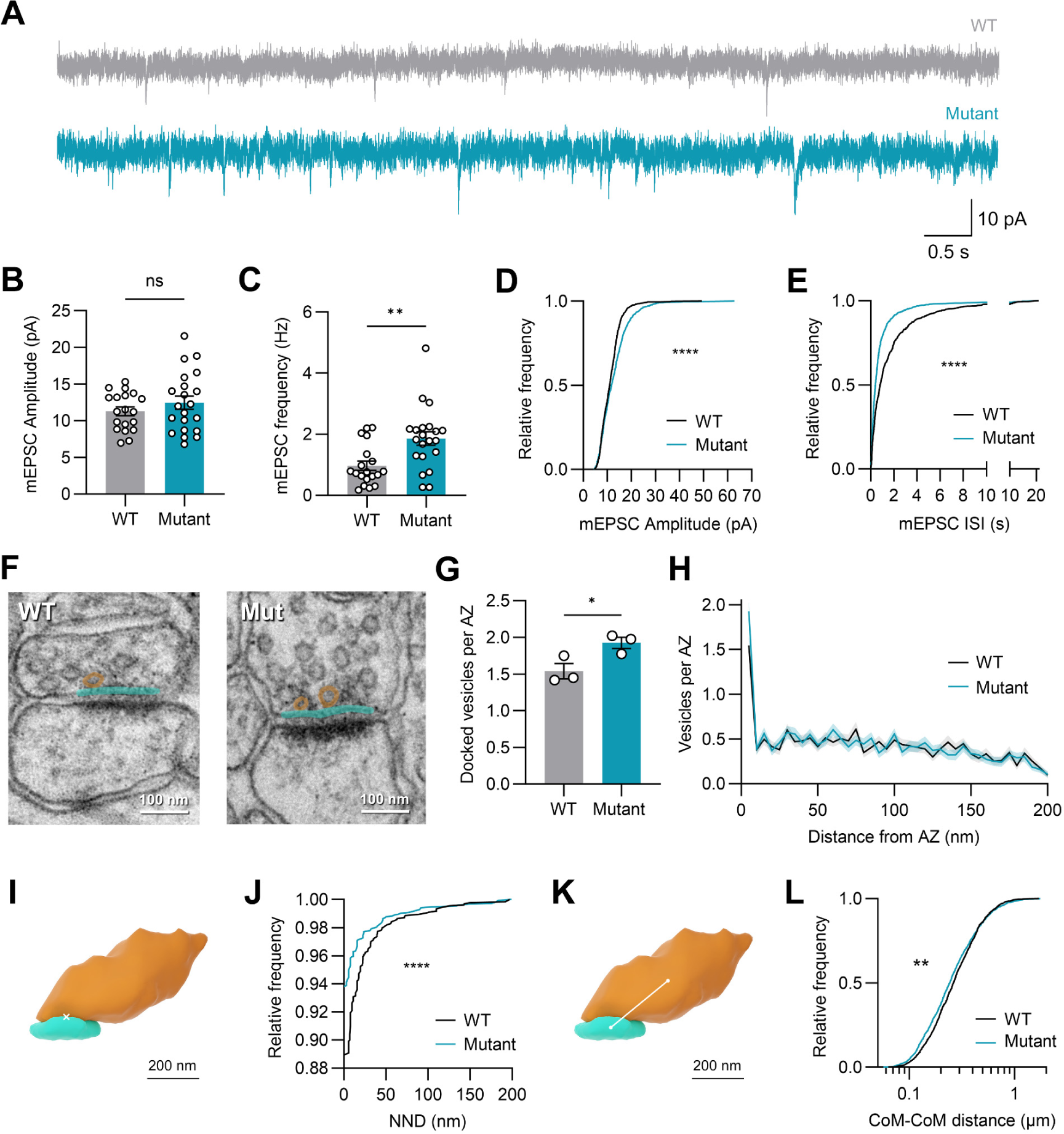
mEPSC frequency and vesicle docking are increased at excitatory synapse of mice with NLGN1 cleavage mutation. **A**. Example traces of mEPSCs for WT and mutant mice. **B**. Mean mEPSC frequency. **C**. Mean mEPSC amplitude. **D**. Cumulative distribution function (CDF) of mEPSC amplitude. **E**. CDF of mEPSC interspike interval (ISI) **F**. Thin-section TEM images of docked synaptic vesicles (orange) at the active zone (AZ, cyan) from both conditions. **G**. Mean docked vesicle number. **H**. Number of vesicles present at increasing distances from the AZ. Bins are 5 nm wide. **I**. 3D rendering of an example synapse (cyan) and paired SVC (orange) showing an NND of 0 nm. **J**. CDF of 3D NNDs from synapses to paired SVCs within less than 200 nm. **K**. 3D rendering showing the CoM distance measure between a synapse and SVC. **L**. CDF of 3D CoM-CoM distances. Error bars represent mean ± S.E.M. * p < 0.05; ** p < 0.01; **** p < 0.0001.

The neuroligin-neurexin complex is known to regulate presynaptic vesicle aggregation^8,36–38^. Docked vesicles contribute to the ready releasable pool^39^; therefore, we measured vesicle docking using thin section TEM (**Figure 3F)**. We found that the frequency of docked vesicles at the AZ in a single section was increased in NLGN1-ΔSD3 mutant mice (mean: WT = 1.54 ± 0.11 vesicles, n = 3 animals; Mutant = 1.92 ± 0.08 vesicles, n = 3 animals, p = 0.0469, t-test; **Figure 3G, H**). We next measured the distance distribution of individual vesicles from the presynaptic membrane and found that individual vesicles that are within 200 nm of the AZ are about 4 nm closer to the membrane in the mutant on median compared to the WT.

We wondered whether the frequency of synapses having vesicles adjacent to the synapse increased in the mutant. Based on vEM reconstructions, we measured the nearest distance of synapses to their paired SVC and found that synapses in NLGN1-ΔSD3 mutant mice more frequently have a vesicle cloud directly adjacent to synapse (89% of WT synapses, n = 1328 synapses; 94% of Mutant synapses, n = 1535 synapses, p < 0.0001, MW test; **Figure 3I, J**). The center of mass distance between synapses and vesicle clouds was also significantly closer by a median of 20 nm (median: WT = 26 nm, IQR = 22 nm, n = 1338 pairs; Mutant = 24 nm, IQR = 22 nm, n = 1577 pairs, p = 0.0012, MW test; **Figure 3K, L**). Taken together, our functional and structural data suggest that release probability and vesicle docking at excitatory synapses in the lateral amygdala of NLGN1-ΔSD3 mutant mice is higher than that of WT mice.

### SVC shape and abundance is unaffected in NLGN1-ΔSD3 mutant mice

As we saw changes to vesicle docking, we wondered whether overall SVC shape or size were affected. We saw no change in the abundance of SVCs in the mutant (mean: WT = 0.401 ± 0.055 clouds/µm^3^, n = 3 animals; Mutant = 0.469 ± 0.005 clouds/µm^3^, n = 3 animals, p = 0.3449, t-test) or the volume fraction occupied by SVCs (mean: WT = 0.020 ± 0.001 µm^3^/µm^3^, n = 3 animals; Mutant = 0.024 ± 0.010 µm^3^, n = 3 animals, p = 0.7206, t-test; **Figure 4A-D**). Despite seeing no difference in overall vesicle content in the tissue, it is possible that the morphology of individual SVCs is impacted by NLGN1 mutation. We therefore compared the volume and shape of SVCs between conditions (**Figure 4E, F**). We saw no difference in the volume of individual SVCs (median: WT = 0.021 µm^3^, IQR = 0.052 µm^3^, n = 1162 SVCs; Mutant = 0.019 µm^3^, IQR = 0.049 µm^3^, n = 1343 SVCs, p = 0.0893; **Figure 4G**). We then measured each vesicle cloud’s sphericity and found no difference between WT and mutant mice (median: WT = 0.620, IQR = 0.1638, n = 1162 SVCs; Mutant = 0.618, IQR = 0.166, n = 1343 SVCs, p = 0.8951, MW test; Figure 4H. Altogether, SVC shape and abundance in the lateral amygdala appear to be unaffected.

**Figure 4.**
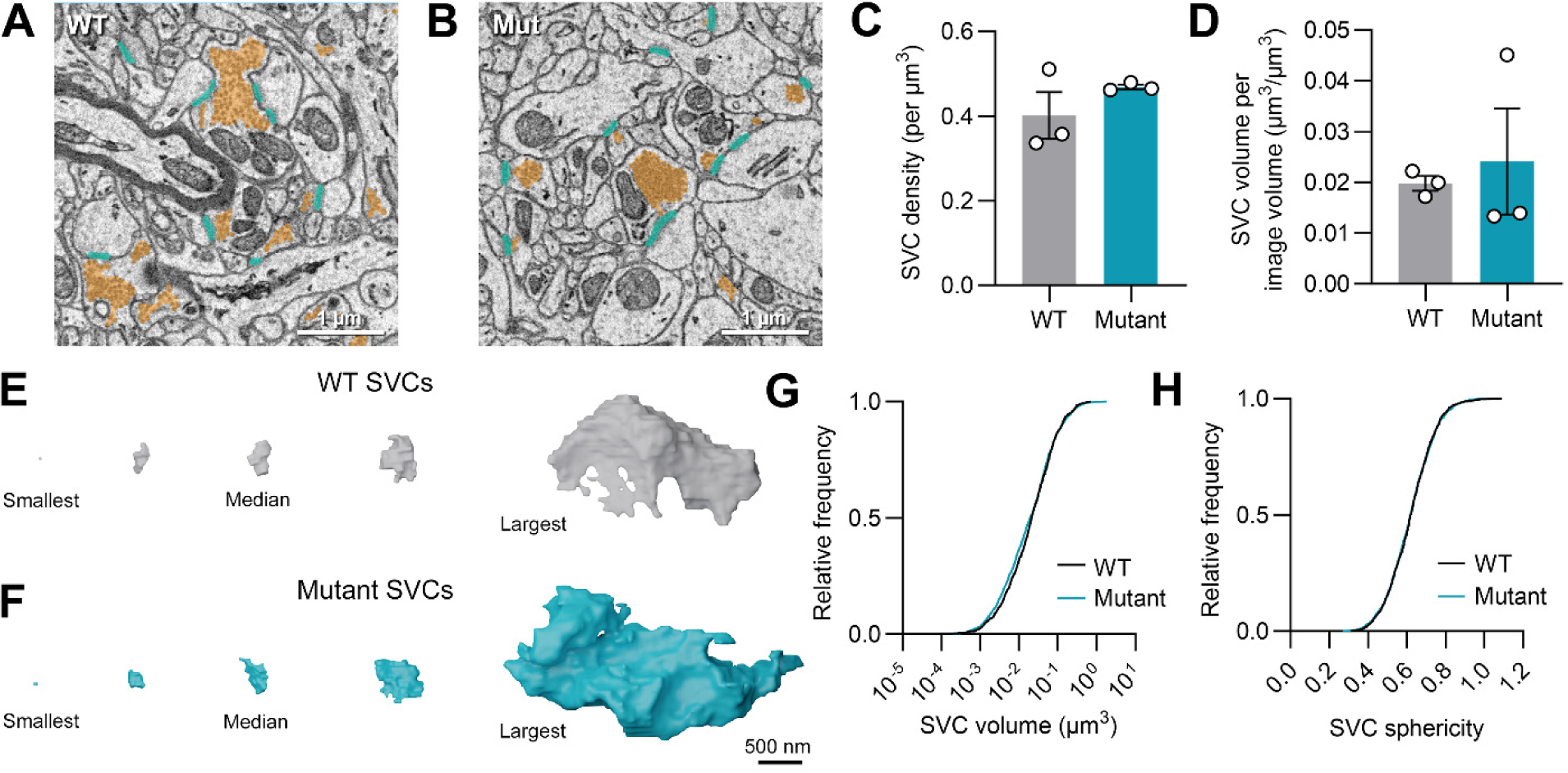
Excitatory SVC abundance and morphology is unaffected by NLGN1 cleavage mutation. **A, B.** SEM images of WT and mutant amygdala, showing synapses (cyan) and vesicles (orange). C. Mean SVC density in the neuropil. **D**. Mean SVC volume per image volume of neuropil. **E**. Gallery of WT SVC reconstructions ordered by size, showing the smallest, median, and largest vesicle clouds by volume. **F**. Gallery of mutant SVC reconstructions. **G**. CDF of SVC volume. **H**. CDF of SVC sphericity. Error bars represent mean ± S.E.M. * p < 0.05.

Our method of pairing synapses to their adjacent SVCs left us with a separate population of unpaired vesicle clouds. These were typically small aggregations of vesicles, sometimes found within presynaptic boutons separate from the SVC, within inter-bouton stretches of axon, or within axo-axonal invaginations. Individual clouds were approximately half the volume in the mutant (median: WT = 0.0064 µm^3^; IQR = 0.010 µm^3^, n = 1013 vesicle clouds; Mutant = 0.0035 µm^3^, IQR = 0.0057 µm^3^, n = 1201 vesicle clouds, p < 0.0001, MW test). In WT animals, the abundance of unpaired clouds was similar, comprising 39-43% of the total population of vesicle clouds. This was much more variable across mutant animals (28-64%).

### Excitatory synapse morphology is altered in NLGN1-ΔSD3 mutant mice, but not abundance

As NLGN1 is a cell adhesion molecule, disrupting its normal function may result in modifications to fine synapse structure, or even result in large scale changes to cell connectivity. To investigate fine synapse structure at high resolution, we performed thin section TEM observation of WT and mutant animals. Then, using our SBF-SEM approach, we measured synapse surface area, complexity, synapse abundance, and volume fraction occupied by synapses.

Modification of synaptic activity has been shown to affect both cleft width and PSD thickness^40,41^. Because activity of mutant synapses is changed, we measured the cleft width as well as PSD thickness using thin section TEM (Figure 5A). Mutant mice had thicker cleft widths (median: WT = 16.89 nm, IQR = 3.68 nm, n = 123 synapses; Mutant = 18.32 nm, IQR = 4.20 nm, n = 124 synapses, p < 0.0001, MW test; Figure 5B), and had considerably thicker PSDs (median: WT = 27.08 nm, IQR = 14.71 nm, n = 123 synapses; Mutant = 39.60 nm, IQR = 16.44 nm, n = 124, p < 0.0001, MW test; Figure 5C).

**Figure 5.**
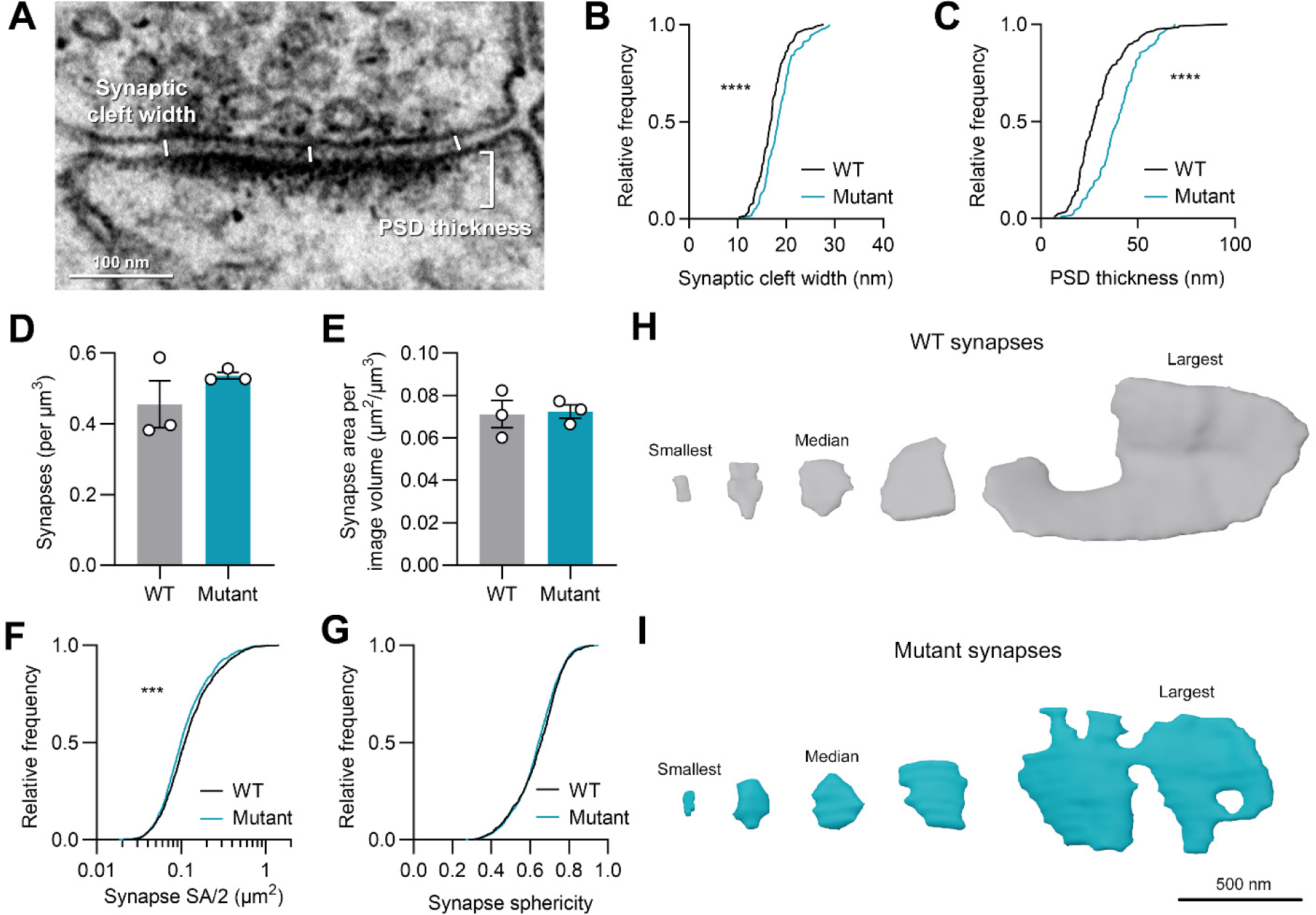
Excitatory synapse morphology is modified by NLGN1 cleavage mutation, but not synapse abundance. **A**. Thin-section TEM image of synapse showing cleft width measurements and PSD thickness. Cleft width was measured as the average of 3 samples. Mean PSD thickness was measured as the PSD cross-sectional area divided by the synapse length. **B**. CDF of mean cleft width. **C**. CDF of mean PSD thickness. **D**. Mean synapse density in the neuropil. **E**. Mean synapse area (surface area/2) per image volume of neuropil. **F**. CDF of synapse area. **G**. CDF of synapse sphericity. **H**. Gallery of WT synapse reconstructions showing the smallest, median, and largest synapses by SA/2. **I**. Gallery of mutant synapse reconstructions. Error bars represent mean ± S.E.M. *** p < 0.001; **** p < 0.0001.

Though SVC abundance was unchanged, it is possible that NLGN1 mutation may alter synapse density in the volume, or synapse size. Neither synapse density in the volume (mean: WT = 0.455 ± 0.066 synapses/µm^3^, n = 3 animals; Mutant = 0.536 ± 0.010 µm^3^, n = 3 animals, p = 0.3445, t-test) nor surface area (SA) per µm^3^ of tissue volume (mean: WT = 0.071 ± 0.006 µm^2^/µm^3^, n = 3 animals; Mutant = 0.072 ± 0.003 µm^2^/µm^3^, n = 3 animals, p = 0.8699, t-test) showed any difference between conditions (Figure 5D**, E**). We then looked at the difference in the distribution of synapse surface areas between conditions and found a very small but statistically significant decrease in synapse area in mutant mice (median: WT = 0.111 µm^2^, IQR = 0.105 µm^2^, n = 1315 synapses; Mutant = 0.099 µm^2^, IQR = 0.095 µm^2^, n = 1540, p < 0.0001, MW test; Figure 5F**, H, I**). Finally, we examined synapse sphericity and saw no difference (median: WT = 0.655, IQR = 0.157, n = 1315 synapses; Mutant = 0.643, IQR = 0.147, n = 1540 synapses, p = 0.0824, MW test; Figure 5G-I). We conclude that synapse morphology is affected by the SD3 mutation, but overall synapse abundance is not.

### Relationships between excitatory synapse and overall vesicle cloud morphology are mildly disrupted in NLGN1-ΔSD3 mutant mice

It is well established that synapse size and vesicle quantity scale strongly together; this relationship is the basis of synaptic strength and assures the efficiency of synaptic transmission^42^. Modification of NLGN1 may shift the balance between synapse size and vesicle accumulation. As we saw an increase in the number of docked vesicles in NLGN1-ΔSD3 mutants using thin section TEM, we next measured and compared the correlation between AZ length and number of docked vesicles for both WT and mutant mice. The correlation of AZ length to number of docked vesicles was significant for both conditions (WT Spearman’s r = 0.4745, n = 123 synapses, p < 0.0001; Mutant Spearman’s r = 0.5564, n = 124 synapses, p < 0.0001), and the two correlations were not different (cocor: Fisher’s z = −0.8673, p = 0.3858; Figure 6A).

**Figure 6.**
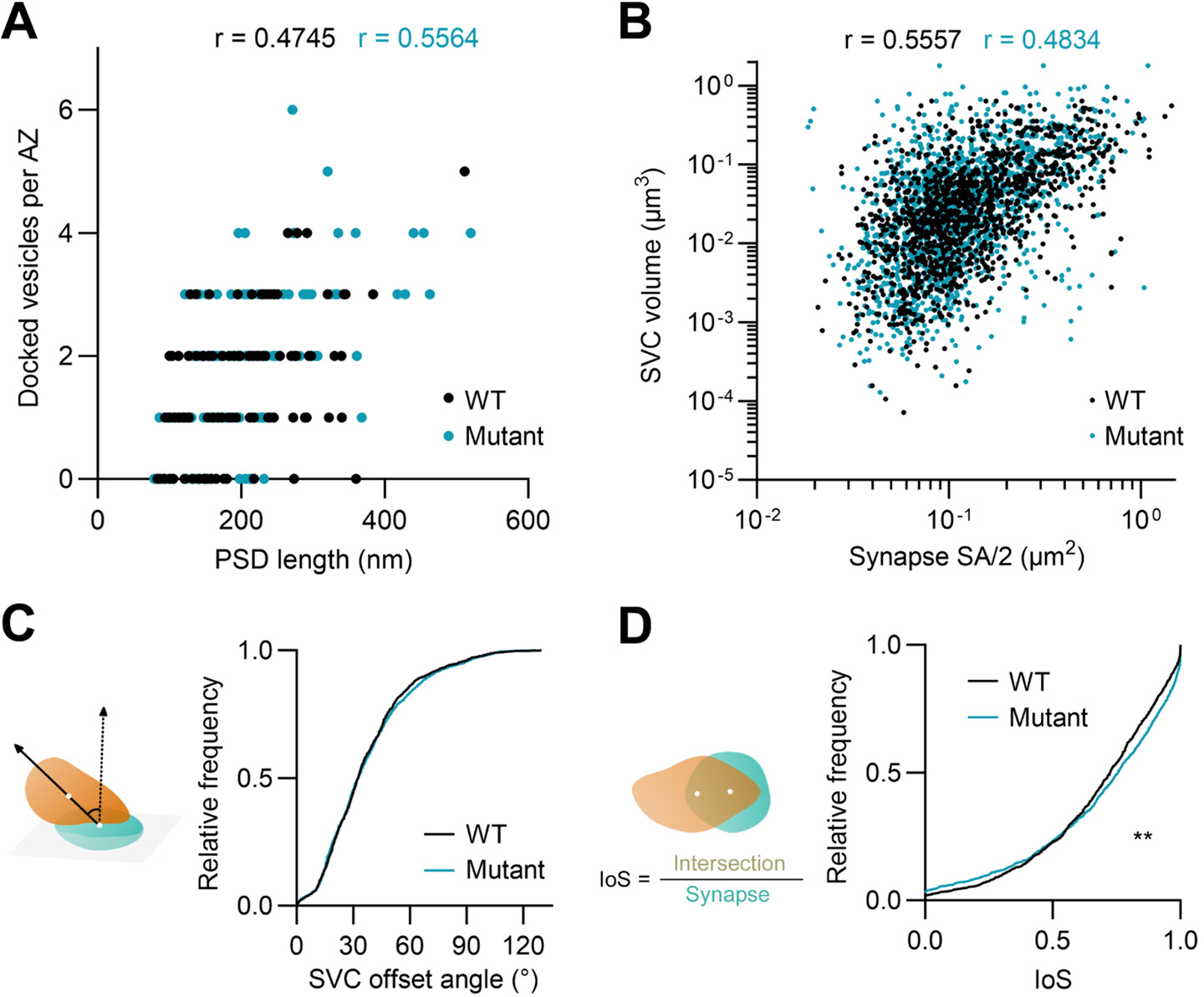
Synapse and vesicle cloud structural and spatial relationships are affected by NLGN1 cleavage mutation. **A.** Correlation between AZ length and number of docked synaptic vesicles. **B.** Correlation between synapse SA/2 and SVC volume. **C.** CDF of SVC offset angle from the synapse. **D.** CDF of SVC intersection relative to the synapse. ** p < 0.01.

We also saw significant correlations between synapse area and SVC volume (WT Spearman’s r = 0.5557, n = 1338 pairs, p < 0.0001; Mutant Spearman’s r = 0.4834, n = 1577 pairs, p < 0.0001; Figure 6B); however, we observed a mild and significant difference in the strength of the correlation in mutant mice compared to WT mice, suggesting that the coupling of vesicle cloud volume to synapse size is disrupted (cocor: Fisher’s z = 2.6656, p = 0.0077).

We next sought to investigate possible misalignment of the synaptic vesicles to the synapse in the NLGN1-ΔSD3 mutants. We applied our analysis pipeline to measure the offset angle between the CoM of the vesicle cloud and CoM of the synapse (Figure 2E), then calculated the IoS (Figure 2F). We saw no difference in the offset angle between WT and mutants (median: WT = 32.50°, IQR = 28.16°, n = 1336 pairs; Mutant = 32.62°, IQR = 30.63°, n = 1577 pairs, p = 0.7551, MW test; Figure 6C), but did see a significant shift in the distribution of the IoS (median: WT = 0.725, IQR = 0.352, n = 1338 pairs; Mutant = 0.755, IQR = 0.400, n = 1577 pairs, p = 0.0087, MW test; Figure 6D). Overall, the 3D geometric relationship between SVCs and their synapses appears to be disrupted in the mutant, causing an increased degree of structural variability in 3D geometry across synapses.

### Prevention of NLGN1 cleavage increases the frequency of incomplete synapse with no vesicle accumulation

While characterizing synapse morphology in both thin section TEM and SBF-SEM, we noticed an increase in the number of incomplete synapses having a postsynaptic compartment but no presynaptic vesicle accumulation. These “empty” synapses were axon-spine interfaces with an asymmetric PSD and rigid membrane apposition, but no presynaptic accumulation of vesicles within at least 200 nm (Figure 7A). Because TEM offers only a single thin slice through a given synapse, we quantified the abundance of incomplete synapses using vEM. Mutants consistently had a greater fraction of synapses without vesicles and a higher density of incomplete synapses per µm^3^ of volume, by roughly a factor of 4 (mean: WT = 0.0085 ± 0.0019 incomplete synapses/µm^3^, n = 3 animals; Mutant = 0.0346 ± 0.0112 incomplete synapses/µm^3^, n = 3 animals, p = 0.143; Figure 7B**, C**). Likely due to our limited sample size and variability between animals, this did not reach statistical significance. Incomplete synapses, pooled across all mutant animals, were significantly smaller than a typical synapse with vesicle accumulation (median: Mutant incomplete synapses = 0.056 µm^2^, IQR = 0.163 µm^2^, n = 52; Mutant typical synapses = 0.103 µm^2^, IQR = 0.098 µm^2^, n = 1524, p < 0.0001, MW test; Figure 7D).

Similar to previous descriptions, most axons across samples had a thin tubular endoplasmic reticulum (ER) that extends the length of the axon, which occasionally forms sheets or cisternae^43^. Intriguingly, at incomplete synapses in both WT and mutant conditions, we often observed close appositions and expansions of the ER lumen (Figure 7E).

**Figure 7.**
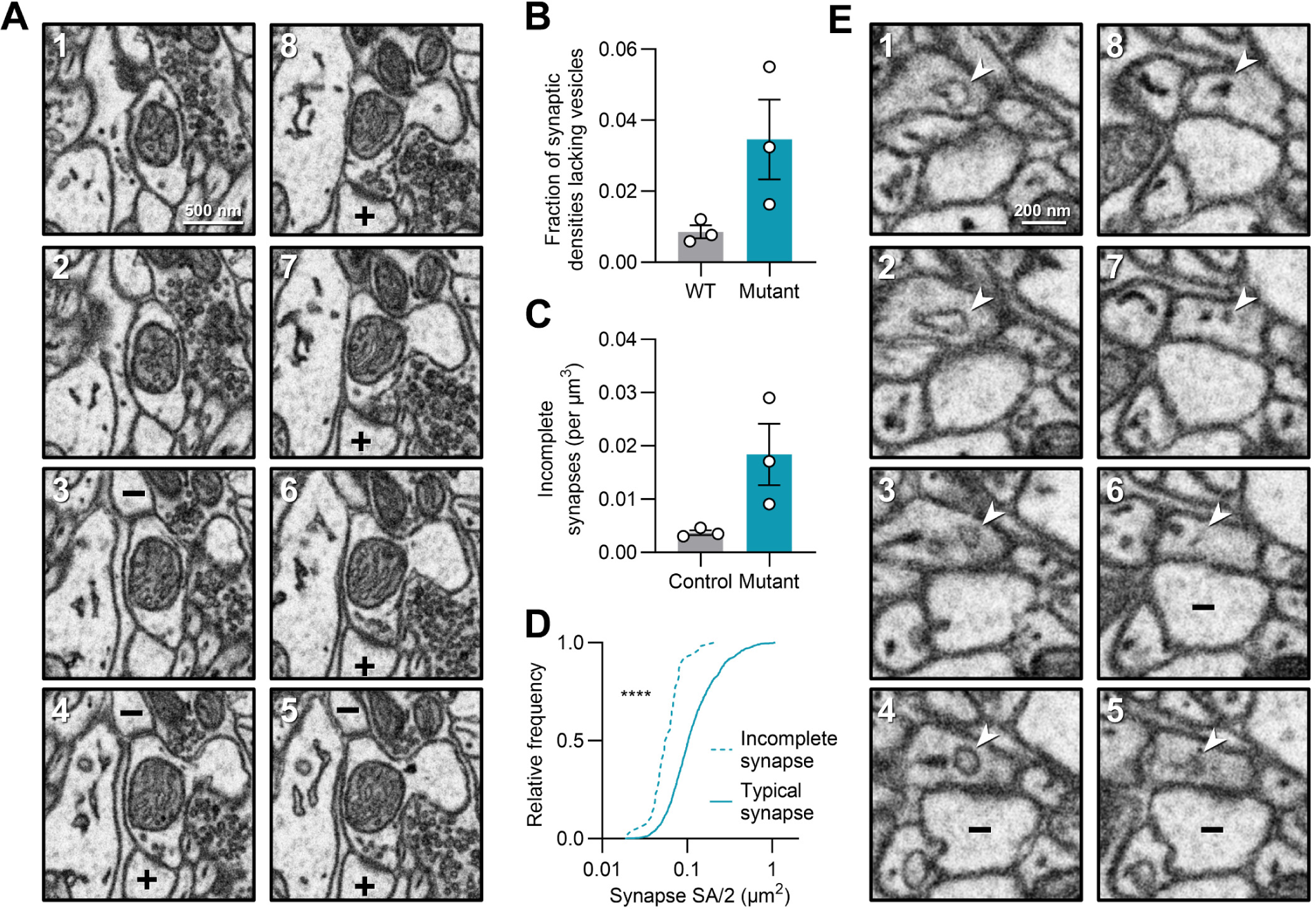
Incomplete synapses are more frequent in the NLGN1 cleavage mutant. **A**. An example SEM image series demonstrating an incomplete synapse lacking vesicle accumulation (top, – in spine) near a typical synapse (bottom, + in spine). **B**. Relative fraction of synapses that lack vesicle accumulation. **C**. Incomplete synapse density in the neuropil. **D**. CDF of incomplete synapse area compared to typical synapses in mutant mice. **E**. An example SEM image series showing an incomplete synapse (– in spine) and expansion of the presynaptic axonal ER lumen. Error bars represent mean ± S.E.M. **** p < 0.0001.

## Methods

### Animals

NLGN1-ΔSD3 knock-in mice were generated by Ingenious targeting laboratory (Ronkonkoma, NY). Gene targeting was performed in iTL IC1 (C57BL/6) ES cells to replace the coding sequences corresponding to amino acids 672-695 of the NLGN1 gene with those encoding amino acids GAAAA. This substitution removes the consensus sequence for metalloprotease mediated cleavage at the extracellular juxta-membrane stalk domain^18^. ES cells were screened and positive clones were microinjected into BALB/c blastocysts and transferred to pseudopregnant female mice. NLGN1(+/+) (WT) were distinguished from NLGN1(ΔSD3/ΔSD3) homozygous mice by genotyping for the presence or absence of the Neomyosin cassette (primer: GGGCGCCCGGTTCTT) and wild-type NLGN1 (AGT AGA GGC AAG TTA ATG TTA AAA CCT AAT TCA). All experiments were performed in accordance with animal welfare laws and approved by the Institutional Committee for Care and Use of Animals at the Max Planck Florida Institute for Neuroscience.

### Electrophysiology

Adult mice (5 WT and 5 Mutant, aged 8-12 weeks) were anesthetized with isoflurane, perfused transcardially with ice cold cutting solution, and then decapitated. The brain was rapidly removed and acute coronal brain slices (350 µm) were prepared in ice cold cutting solution containing (in mM): 124 choline chloride, 26 NaHCO3, 2.5 KCl, 3.3 MgCl2, 1.2 NaH2PO4, 10 glucose, and 0.5 CaCl2 (osmolarity adjusted to 300 mOsm). Slices were allowed to recover for 30 min at 32° C then were stored at room temperature in artificial cerebrospinal fluid (ACSF) containing (in mM): 124 NaCl, 26 NaHCO3, 3 KCl, 1.25 NaH2PO4, 20 glucose, 1 MgCl2, 2 CaCl2, 5 sodium ascorbate, 3 sodium pyruvate, and 2 thiourea (300 mOsm). All solutions were continuously bubbled with 95% O2/5% CO2. Slices containing the lateral amygdala were transferred to a submersion recording chamber, superfused with oxygenated ACSF at a speed of 1–2 ml/min. Whole-cell patch-clamp recordings were performed using pipettes pulled from borosilicate glass capillaries (BF150-110-10, Sutter Instruments) with resistances of 3–6 MW. Lateral amygdala principal neurons were visualized for whole-cell recording with infrared differential contrast optics on an Olympus BX51WI microscope. Data was acquired using Clampex 10.7 software with an Axoclamp 700B amplifier and digitized at 20 kHz with a Digidata 1440A interface and filtered at 2 kHz (Molecular Devices). Recordings where access was greater than 30MW or that changed by >20% were discarded. AMPA mediated mEPSCs were recorded in voltage clamp mode for 3 min at −70 mV using a cesium gluconate based internal solution containing (in mM): 140 Cs-gluconate, 8 NaCl, 10 HEPES, 0.2 EGTA, 2 MgATP, and 0.2 Na2 GTP (285 mOsm, pH 7.25). Action potentials were blocked with 2 µM TTX-citrate, NMDA currents with 50 µM APV and inhibitory currents with 20 µM Picrotoxin. mEPSCs were detected and quantified using the template detection algorithm in Clampfit (Molecular Devices).

### Perfusion fixation

Tissue slices for EM were prepared independently of acute slices used for electrophysiology experiments. Adult mice aged 13-15 (SBF-SEM) or 17 (TEM) weeks were anesthetized and perfused transcardially with 0.9% NaCl in water, followed by a fixative containing 2% paraformaldehyde and 2.5% glutaraldehyde in 0.15 M cacodylate buffer (CB), pH 7.4, containing 2 mM CaCl2. Brains were extracted and postfixed in the same fixative overnight, then washed in CB. Brain tissue was sliced at a thickness of 100 μm using a Leica VT 1200S vibratome in 0.1 M CB. For SBF-SEM, the lateral amygdala was trimmed prior to sample preparation for EM.

### TEM sample processing and imaging

Slices were incubated in 1% aqueous Osmium tetroxide (OsO4) for 1 h, rinsed with distilled water, incubated in 1% aqueous uranyl acetate in the dark for 1 h at 4°C, then rinsed again with distilled water. Slices were dehydrated in a graded ethanol series (30, 50, 70, 90, 100%), followed by 1:1 ethanol to acetone, 100% acetone, 1:1 acetone to propylene oxide (PO), and 100% PO, each 2 × 5 min. Tissue was infiltrated using 3:1 PO to Durcupan resin (Sigma-Aldrich) for 2 h, 1:1 PO to resin for 2 h, and 1:3 PO to resin overnight, then flat embedded in 100% resin on a glass slide and covered with an Aclar sheet at 60°C for 2 d. The lateral amygdala was trimmed, mounted on a resin block with cyanoacrylate glue (Krazy Glue, Elmer’s Products), and sectioned at 40 nm using a UC7 ultra-microtome (Leica Microsystems). Sections were counterstained with uranyl acetate and lead citrate, and examined in a Tecnai G2 Spirit BioTwin transmission electron microscope (ThermoFisher Scientific) at 100 kV acceleration voltage. Images were taken with a Veleta CCD camera (Olympus) operated by TIA software (ThermoFisher Scientific). Images used for quantification were taken at 60,000 × magnification.

### TEM image analysis

AZs were identified by the following criteria: (1) rigid opposed presynaptic and postsynaptic membrane, (2) SVs clustering at the membrane, and (3) an asymmetric postsynaptic density. All TEM data were analyzed using the Fiji software^44^. Vesicle distance to the AZ was calculated using a 32-bit Euclidean distance map generated from the AZ for all vesicles within 200 nm of the presynaptic membrane. Vesicles within 5 nm of the AZ were considered to be docked^45–47^. Cleft width was measured by calculating the average of three widths (two at the ends, one in the center, see Figure 5A) between the outer cell membranes. PSD thickness was calculated by using the polygon tool to manually demarcate the PSD based on electron density to get an area, which was then divided by the PSD length to obtain an average PSD thickness.

### SBF-SEM sample processing and imaging

Following trimming, tissue pieces were incubated in an aqueous reduced osmium solution (2% OsO4 and 1.5% potassium ferrocyanide) for 1 h on ice and in the dark. Tissue was washed with water, which was repeated between each of the following steps. Tissue was then reacted with 1% aqueous thiocarbohydrazide heated to 60°C for 20 min, 2% aqueous osmium tetroxide for 30 min, then incubated in 1% uranyl acetate overnight at 4°C. The following day, samples were reacted with Walton’s lead aspartate for 30 min at 60°C. Dehydration, resin infiltration, and embedding were performed as described in the previous TEM sample processing section. Once placed on a resin block, one side was exposed using an ultra-microtome (UC7, Leica), then flipped to be remounted to a metal pin with conductive silver epoxy (CircuitWorks, Chemtronics). The new surface was then exposed and sputter-coated with a 5 nm layer of platinum. Samples were then sectioned and imaged using 3View and Digital Micrograph (Gatan Microscopy Suite) installed on a Gemini SEM300 (Carl Zeiss Microscopy) equipped with an OnPoint BSE detector (Gatan). The detector magnification was calibrated before imaging within SmartSEM imaging software (Carl Zeiss Microscopy) and Digital Micrograph with a 500 nm cross line grating standard. Imaging conditions were set on a sample-by-sample basis. Imaging was performed at 1.4-2.2 kV accelerating voltage, 20-30 μm aperture, working distance of ∼5 mm, 0.5-0.8 μs pixel dwell time, 4-10 nm pixel size, N2 focal charge compensation setting of 0-20%, and 45-60 nm section thickness. Serial images were aligned using the Digital Micrograph software and converted to TIFF format, or exported as TIFFs to TrakEM2 and aligned using Scale-Invariant Feature Transform image alignment with linear feature correspondences and rigid transformation^48^.

### Image processing, deep learning network segmentation, and proofreading

Aligned images were imported into ORS Dragonfly (ver. 2022.2) for processing, segmentation, and analysis. Volumes corresponding to 1000 µm^3^ of the lateral amygdala (approximately 10 × 10 × 10 µm) of WT (n = 3) and Mutant (n = 3) animals were cropped to standardize volume calculations. Initially, a 2.5D U-Net++ architecture was used, then a Sensor3D architecture as performance was observed to be higher (Figure 1D). The Sensor3D architecture was originally created to be used on 3D computed tomography (CT) scans^49^; nonetheless, it also performed well on 3D EM data (Figure 1C). The first models were trained starting from random weights, on a small set of manually segmented synapses and vesicle clouds (∼20 complete 3D segmentations of each). As more image data was collected, networks were retrained on the new datasets, as slight differences in image quality and contrast prevented direct application of the same network. Some synapses were sectioned *en face* such that no cleft was clearly visible, in which case the synapse was labeled based on the shape of the PSD. Since NLGN1 is found at excitatory synapses, and asymmetric synapses are regarded as excitatory^50^, we only included asymmetric synapses in our training data and analysis. Vesicle clouds were defined as being at least 3 vesicles within approximately 100 nm of each other. Prior to training, a gaussian blur was applied to each dataset to improve network performance. We extracted vesicles as clouds because individual vesicles are difficult to extract with this technique, which is limited by section thickness, signal-to-noise ratio, and XY resolution. Once an output was generated, we began the process of proofreading. The output was then filtered to remove objects (i.e. synapses and vesicle clouds) that intersected with the edges of the image volume, as well as those below a 100 voxel volume threshold. Segmentations were proofread by two human annotators, and an expert annotator with over 1000 hours of segmentation experience then manually corrected the resulting segmentations. Proofreading time varied depending on network performance and image quality, but ranged from 9-19.5 hours per dataset as shown in Figure 1E. Proofreading was mostly necessary to remove the false positive segmentation of mitochondria and ER processes, the latter often having a vesicle-like appearance in one or more sections. These were differentiable from synaptic vesicles as they could be traced through several sections. Object islands were separated using the Multi-ROI tool in Dragonfly, and statistics were generated for each isolated object. For neuropil volume-density calculations, volume taken up by portions of somata, large myelinated axons, and large proximal dendrites were subtracted from the volumes. Vesicle and synapse sphericity was measured by finding the SA of a sphere with equal volume, then dividing the sphere’s SA by the SA of the vesicle cloud or synapse.

### Vesicle cloud and synapse analysis pipeline

We created a custom set of plugins designed for the ORS Dragonfly software for mesh export, mesh based synapse-vesicle NND pairing, and vesicle offset and intersection calculation. Installation instructions and code are openly available on Github at https://github.com/mpfi-dsp/Syn2Ves-User. Dragonfly ORS is free for non-commercial use.

Following proofreading, meshes of synapses and vesicle clouds were batch exported from Dragonfly, along with a CSV of corresponding mesh indices ordered by size. For each synapse, all vesicle clouds with a CoM within 2 µm of the synapse CoM were surveyed. For each vertex on the synapse mesh, the NND to a vertex on the vesicle clouds within the window was calculated. The cloud with the shortest NND was labeled as the correct pairing for that synapse. In this way, each synapse has exactly one paired vesicle cloud, unless multiple vesicle clouds were 0 nm from one synapse. Mesh pairing accuracy of synapses to vesicle clouds was manually, visually assessed for a sampling of one-third of the data collected, and was verified to be 98.8%–100% accurate across animals.

Vesicle-synapse overlap was measured from a perspective that captures the synapse’s maximum surface area (SA) from an orthographic projection. Using the PyVista library in Python, we acquired 2D projections of the synapse mesh, then imposed a projection of vesicle cloud. Intersection over synapse (IoS) area was calculated as:

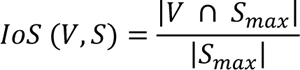

where *V* represents the projection of the vesicle cloud, *S_max_* represents the maximum projection of the synapse, |*S_max_*| represents the area of the synapse, and |*V* ∩ *S_max_*| represents the area of intersection between the two projections.

The offset angle was calculated as the angle (*θ*) between two 3D vectors A and B using the dot product and magnitudes of the vectors:

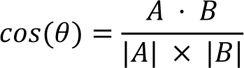

where *A* ⋅ *B* represents the dot product of the two vectors, and |*A*| and |*B*| are the magnitudes of those vectors. One vector describes the line between the synapse and SVC CoMs, the other describes the line between the synapse CoM and the position from which a virtual camera captures the synapse’s maximum SA projection (see Figure 2).

### Statistics

Two-tailed Student’s t-tests with Welch’s correction were applied for all comparisons of means. For non-normally distributed data, including data shown as cumulative distributions, two-tailed Mann-Whitney *U* tests (MW) were applied. For all statistical tests, significance was set at α = 0.05. Values reported in the text are means ± standard error (S.E.M) for t-tests and medians with interquartile range (IQR) for MW tests. To measure the strength of monotonic relationships, Spearman’s correlation coefficient was calculated. To statistically compare correlations, the cocor package^51^ was used in RStudio (ver. 2023.03.0+386). All other statistics were calculated using GraphPad Prism (ver. 10).

## Discussion

In the present work, we established a novel analysis strategy by combining mEPSC recordings, thin-section TEM, and vEM analysis to investigate how cleavage of NLGN1 affects synaptic function and structure in the lateral amygdala. We developed and utilized a pipeline for extraction of large quantities of synapses and vesicle clouds from vEM data, followed by pairing of vesicle clouds to their release sites, and characterization of their geometric relationships. Through mutation of the juxtamembrane stalk domain of NLGN1, we show that prevention of cleavage leads to a general strengthening of synaptic transmission, while also resulting in an increased frequency of incomplete synapses. Altogether, by applying a multi-faceted approach, we demonstrate that NLGN1 cleavage plays a role in synaptic strength and possibly synaptic turnover in the lateral amygdala.

### Large-Scale 3D Structural Quantification

While 3D analysis techniques are growing in commonality, vEM investigation comes with many challenges. Image acquisition of brain tissues is largely automated, leading to the collection of massive datasets—but analysis remains very labor intensive. Manual segmentation of hundreds to thousands of synapses and related features can require months to years of manual work^26,52^, and even once segmentation is completed, manual quantification of such a number of features is impractical. We used U-Net++ and Sensor3D networks to extract thousands of features with minimal manual labor, and showed that Sensor3D networks are high performing, despite being designed for CT image segmentation^49^. Still, alternative architectures such as region-based convolutional neural networks and similar algorithms are being developed constantly, and are shown to be highly effective at segmenting synaptic features^29–31^.

Our pipeline differs from others in that it is the first attempt to focus on characterizing the structural relationships between thousands of synapses and vesicle clouds. Vesicle aggregation and distance to the synapse is a critical readout of synapse function^53,54^, and the strong correlation between synapse size and vesicle accumulation serves as a readout of synaptic strength^42^. Therefore, our approach becomes universally applicable when studying the effects synaptic protein perturbation. To showcase our pipeline, we induced a mutation in NLGN1 to holistically characterize the effects cleavage at the SD3 region has on both ultrastructure and abundance of synapses within the neuropil.

### Functional and Structural Effects of NLGN1 Cleavage in the Lateral Amygdala

Functional studies of NLGN1 cleavage by the proteases MMP9 and ADAM10 have been primarily performed in dissociated neurons^18,19^. Although cleaved fragments have been observed by immunoblot in cerebellar, cortical, and hippocampal tissue fractions following in vivo cleavage^18^, characterization of the functional and structural effects of normal NLGN1 cleavage in brain tissue has yet to be tackled. Our electrophysiological data show a substantial increase in the frequency of miniature release events, which can be structurally explained by our EM observation of increased vesicle docking and shorter distance of vesicles to the presynaptic membrane. Additionally, by investigating a large sample of synapses in tissue volumes, we found that the incidence of SVCs in apposition to the synapse increased when cleavage is blocked. While vesicle clouds are shifted towards the synapse, SVC morphology is largely unchanged, as evidenced by a lack of change to SVC volume or sphericity. These data suggest transsynaptic control of vesicle aggregation via NLGN1 cleavage mechanisms is mainly limited to docking, and does not change the total number of vesicles at the presynaptic terminal. This is supported by evidence that links NLGN1 and neurexin-1β activity directly to the ready releasable pool^55,56^.

While our findings demonstrate an increase in vesicle release, it remains unclear exactly why this is the case. One explanation is that blocking cleavage results in a build-up of NLGN1 at the synapse at higher-than-normal levels. Increased levels of NLGN1 may attract neurexin-1β, which in turn recruits synaptic vesicles^9^. The neurexin-neuroligin complex binds and recruits PSD95 to the synapse^7,10^, providing a possible explanation of the thicker PSDs we observed in NLGN1-ΔSD3 mutants.

As NLGN1 is known to be involved in synapse formation and maturation, it was possible that mutants may have an increased density of excitatory synapses per unit volume or the structural relationship between synapses and SVCs. Overexpression of NLGN1 in the hippocampus increases spine and synapse density, reserve pool size, and excitation^57^. In our study, synapse abundance in the neuropil was variable among animals and was not significantly different in the mutant. Interestingly, the relationship between synapse area and SVC volume was slightly weaker in the mutant and significantly different from WT. Despite this, the correlation was still significant, suggesting only a mild miscoordination. A change in the offset of the vesicle cloud positioning relative to synapse might occur if the mechanisms that control vesicle aggregation at the synapse are affected. We observed no shift in the vesicle cloud offset angle, and the intersection between the synapse and vesicle cloud projection was only slightly different. Overall, our findings suggest NLGN1 cleavage does not affect synapse or vesicle abundance in the neuropil, and only mildly disrupts structural relationships, though this seemingly does not impact vesicle release.

Our analysis pipeline allowed us to separate vesicle clouds into two populations—those in apposition to a synapse and those isolated from one. This second population of unpaired vesicles, sometimes referred to as orphan, superpool, or mobile vesicles, potentially serve as mobile reserves that travel along the axon between synapses^58–64^. While their overall abundance was highly variable between animals, we observed a significant and considerable decrease in the size of individual unpaired vesicle clouds in mutant animals. As this mobile pool has been reported to replenish the recycling pool^59^, it is possible that increased vesicle release may result in depletion of unpaired vesicles in the mutant.

We observed a four-fold increase in the frequency of incomplete synapses that do not have synaptic vesicle accumulation in mutant tissues. We hypothesize that these are remnants of synapses that are undergoing disassembly. In the mutant, more synapses may persist at this stage, as NLGN1 cleavage is prevented and synaptic turnover is slowed (Figure 8). Still, as pruning occurs throughout the life of an animal, why is it that we do not see an even greater frequency of empty synapses if cleavage is completely blocked? It is possible that some degree of cleavage is still occurring. In addition to the n-terminal ectodomain cleavage by MMP9 and ADAM10 at the stalk domain, the intracellular c-terminal fragment is cleaved by presenilin/γ-secretase^19^. Cleavage of neurexin-1β by ADAM10 may also allow for some degree of synaptic turnover^65^. Therefore, postsynaptic intracellular NLGN1 cleavage or presynaptic neurexin-1β cleavage may provide pathways for synapse disassembly, but blocking NLGN1 ectodomain cleavage may slow the process. Finally, we observed lumenal expansions of ER tubules at incomplete synapses. Presynaptic smooth ER are critical for lipid biosynthesis and protein export^66^, but the function of these particular structures is unclear as we found no previous description of them in our review of the literature.

**Figure 8.**
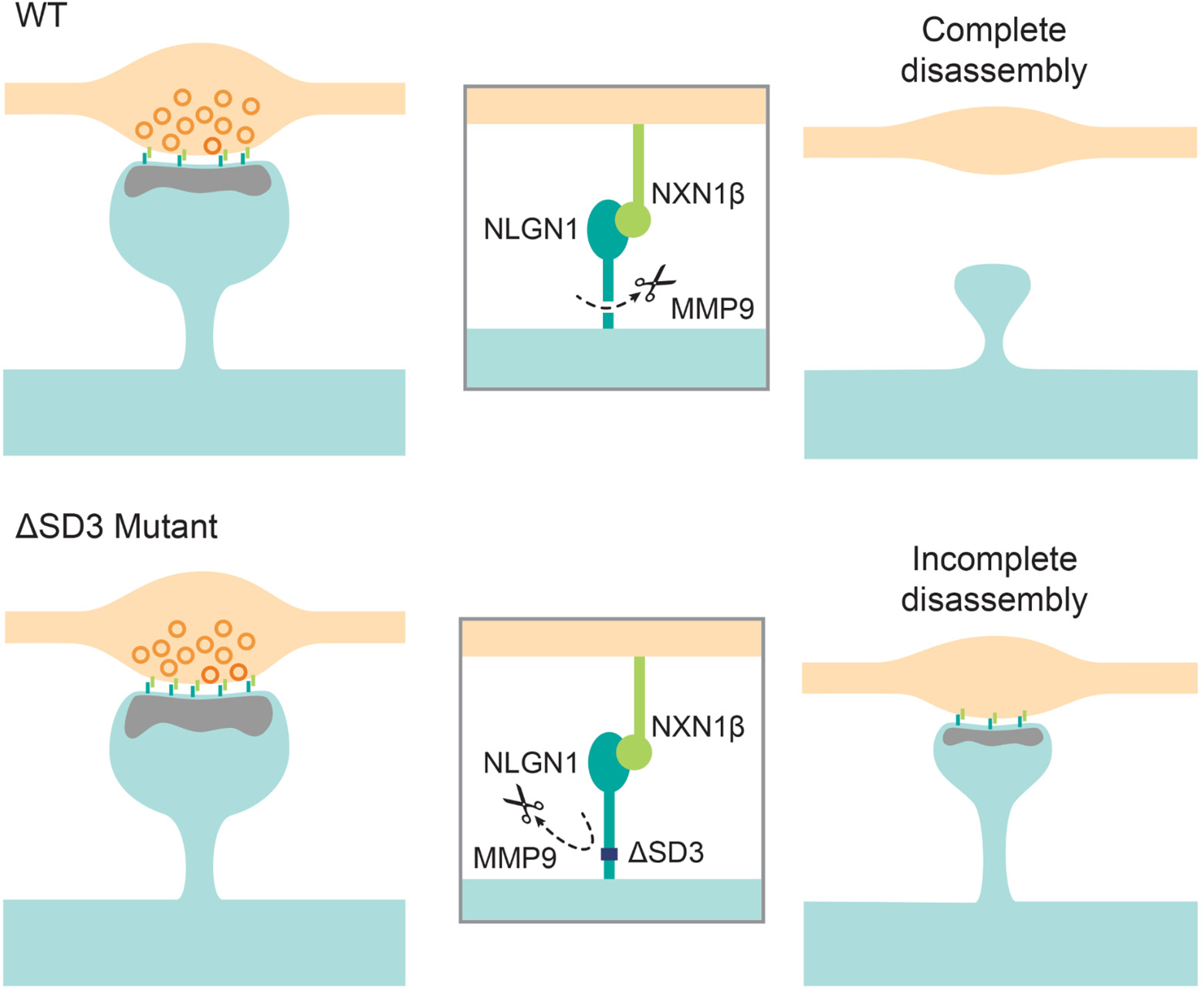
A putative model of the effects of NLGN1-ΔSD3 mutation on synapse structure and function. Postsynaptic NLGN1 binds transsynaptically to neurexin-1β (NXN1β). In WT mice, NLGN1 undergoes activity dependent cleavage to depress synaptic transmission. In the ultimate case, synapses are able to be disassembled through pruning mechanisms. In the ΔSD3 mutant, NLGN1 is unable to undergo ectodomain cleavage, resulting in a buildup of NLGN1 at the synapse and increased recruitment of PSD95 and neurexin-1β and subsequent increase of vesicle docking and release. Synapses that are set to be disassembled are done so at a slower rate or are unable to be completely broken down, resulting in incomplete synapses that lack axonal vesicle accumulation but still adhere spine and axon compartments.

Our study applied a multi-faceted analysis in order to reveal new insights for synaptic organization in brain tissue, and has the potential to provide an analysis standard for characterizing synapse and vesicle structure in animal models having deficiency or mutations in synaptic machinery.

## Author Contributions

M.M.B, N.K., and C.I.T. conceived the project. M.D.E provided the NLGN1-ΔSD3 knock-in mice. M.M.B, A.C., and B.U. performed electrophysiological experiments. C.I.T., M.C.M, J.R.A., and D.G-G. performed electron microscopy imaging. C.I.T., M.C.M., and J.R.A. performed network training and segmentations. C.I.T. performed TEM and SEM data analysis. A.A., J.R.A., and C.I.T. developed the 3D analysis pipeline. C.I.T. constructed figures and drafted the manuscript with input from the authors. All authors contributed to the article and approved the submitted version.

## Data Availability

Dragonfly plugins for mesh export, mesh based synapse-vesicle NND pairing, and vesicle offset and intersection calculation are openly available on Github at https://github.com/mpfi-dsp/Syn2Ves-User.

## References

1. Song, J.-Y., Ichtchenko, K., Südhof, T. C. & Brose, N. Neuroligin 1 is a postsynaptic cell-adhesion molecule of excitatory synapses. Proceedings of the National Academy of Sciences 96, 1100–1105 (1999).

2. Craig, A. M. & Kang, Y. Neurexin–neuroligin signaling in synapse development. Curr Opin Neurobiol 17, 43 (2007).

3. Südhof, T. C. Neuroligins and neurexins link synaptic function to cognitive disease. Nature 455, 903–911 (2008).

4. Nakanishi, M. et al. Functional significance of rare neuroligin 1 variants found in autism. PLOS Genetics 13, e1006940 (2017).

5. Arias-Aragón, F. et al. A Neuroligin-1 mutation associated with Alzheimer’s disease produces memory and age-dependent impairments in hippocampal plasticity. iScience 26, 106868 (2023).

6. Ichtchenko, K., Nguyen, T. & Südhof, T. C. Structures, Alternative Splicing, and Neurexin Binding of Multiple Neuroligins (*). Journal of Biological Chemistry 271, 2676–2682 (1996).

7. Irie, M. et al. Binding of Neuroligins to PSD-95. Science 277, 1511–1515 (1997).

8. Scheiffele, P., Fan, J., Choih, J., Fetter, R. & Serafini, T. Neuroligin Expressed in Nonneuronal Cells Triggers Presynaptic Development in Contacting Axons. Cell 101, 657–669 (2000).

9. Dean, C. et al. Neurexin mediates the assembly of presynaptic terminals. Nat Neurosci 6, 708– 716 (2003).

10. Graf, E. R., Zhang, X., Jin, S.-X., Linhoff, M. W. & Craig, A. M. Neurexins Induce Differentiation of GABA and Glutamate Postsynaptic Specializations via Neuroligins. Cell 119, 1013–1026 (2004).

11. Chubykin, A. A. et al. Activity-Dependent Validation of Excitatory versus Inhibitory Synapses by Neuroligin-1 versus Neuroligin-2. Neuron 54, 919–931 (2007).

12. Mondin, M. et al. Neurexin-Neuroligin Adhesions Capture Surface-Diffusing AMPA Receptors through PSD-95 Scaffolds. J. Neurosci. 31, 13500–13515 (2011).

13. Budreck, E. C. et al. Neuroligin-1 controls synaptic abundance of NMDA-type glutamate receptors through extracellular coupling. Proceedings of the National Academy of Sciences 110, 725–730 (2013).

14. Giannone, G. et al. Neurexin-1β Binding to Neuroligin-1 Triggers the Preferential Recruitment of PSD-95 versus Gephyrin through Tyrosine Phosphorylation of Neuroligin-1. Cell Reports 3, 1996–2007 (2013).

15. Bemben, M. A., Shipman, S. L., Nicoll, R. A. & Roche, K. W. The cellular and molecular landscape of neuroligins. Trends in Neurosciences 38, 496–505 (2015).

16. Haas, K. T. et al. Pre-post synaptic alignment through neuroligin-1 tunes synaptic transmission efficiency. eLife 7, e31755 (2018).

17. Letellier, M. et al. A unique intracellular tyrosine in neuroligin-1 regulates AMPA receptor recruitment during synapse differentiation and potentiation. Nat Commun 9, 3979 (2018).

18. Peixoto, R. T. et al. Transsynaptic Signaling by Activity-Dependent Cleavage of Neuroligin-1. Neuron 76, 396–409 (2012).

19. Suzuki, K. et al. Activity-Dependent Proteolytic Cleavage of Neuroligin-1. Neuron 76, 410–422 (2012).

20. Bozdagi, O., Nagy, V., Kwei, K. T. & Huntley, G. W. In Vivo Roles for Matrix Metalloproteinase-9 in Mature Hippocampal Synaptic Physiology and Plasticity. Journal of Neurophysiology 98, 334–344 (2007).

21. Nagy, V. et al. Matrix Metalloproteinase-9 Is Required for Hippocampal Late-Phase Long-Term Potentiation and Memory. J. Neurosci. 26, 1923–1934 (2006).

22. Wang, X. et al. Extracellular proteolysis by matrix metalloproteinase-9 drives dendritic spine enlargement and long-term potentiation coordinately. Proc. Natl. Acad. Sci. U.S.A. 105, 19520–19525 (2008).

23. Michaluk, P. et al. Influence of matrix metalloproteinase MMP-9 on dendritic spine morphology. Journal of Cell Science 124, 3369–3380 (2011).

24. De Stefano, M. E. & Herrero, M. T. The multifaceted role of metalloproteinases in physiological and pathological conditions in embryonic and adult brains. Progress in Neurobiology 155, 36–56 (2017).

25. Nagappan-Chettiar, S., Johnson-Venkatesh, E. M. & Umemori, H. Activity-dependent proteolytic cleavage of cell adhesion molecules regulates excitatory synaptic development and function. Neuroscience Research 116, 60–69 (2017).

26. Kornfeld, J. & Denk, W. Progress and remaining challenges in high-throughput volume electron microscopy. Current Opinion in Neurobiology 50, 261–267 (2018).

27. Peddie, C. J. et al. Volume electron microscopy. Nat Rev Methods Primers 2, 1–23 (2022).

28. Steinkellner, T. et al. Genetic Probe for Visualizing Glutamatergic Synapses and Vesicles by 3D Electron Microscopy. ACS Chem. Neurosci. 12, 626–639 (2021).

29. Liu, J. et al. Fear memory-associated synaptic and mitochondrial changes revealed by deep learning-based processing of electron microscopy data. Cell Reports 40, 111151 (2022).

30. Su, F. et al. Deep learning-based synapse counting and synaptic ultrastructure analysis of electron microscopy images. Journal of Neuroscience Methods 384, 109750 (2023).

31. Aswath, A., Alsahaf, A., Giepmans, B. N. G. & Azzopardi, G. Segmentation in large-scale cellular electron microscopy with deep learning: A literature survey. Medical Image Analysis 89, 102920 (2023).

32. Kim, J. et al. Neuroligin-1 is required for normal expression of LTP and associative fear memory in the amygdala of adult animals. Proceedings of the National Academy of Sciences 105, 9087–9092 (2008).

33. Jung, S.-Y. et al. Input-specific synaptic plasticity in the amygdala is regulated by neuroligin-1 via postsynaptic NMDA receptors. Proceedings of the National Academy of Sciences 107, 4710–4715 (2010).

34. Avesta, A. et al. Comparing 3D, 2.5D, and 2D Approaches to Brain Image Auto-Segmentation. Bioengineering 10, 181 (2023).

35. Silva, M., Tran, V. & Marty, A. Calcium-dependent docking of synaptic vesicles. Trends in Neurosciences 44, 579–592 (2021).

36. Futai, K. et al. Retrograde modulation of presynaptic release probability through signaling mediated by PSD-95–neuroligin. Nat Neurosci 10, 186–195 (2007).

37. Stan, A. et al. Essential cooperation of N-cadherin and neuroligin-1 in the transsynaptic control of vesicle accumulation. Proceedings of the National Academy of Sciences 107, 11116–11121 (2010).

38. Rui, M. et al. The neuronal protein Neurexin directly interacts with the Scribble–Pix complex to stimulate F-actin assembly for synaptic vesicle clustering. Journal of Biological Chemistry 292, 14334– 14348 (2017).

39. Kaeser, P. S. & Regehr, W. G. The readily releasable pool of synaptic vesicles. Current Opinion in Neurobiology 43, 63–70 (2017).

40. Dosemeci, A. et al. Glutamate-induced transient modification of the postsynaptic density. Proceedings of the National Academy of Sciences 98, 10428–10432 (2001).

41. Glebov, O. O., Cox, S., Humphreys, L. & Burrone, J. Neuronal activity controls transsynaptic geometry. Sci Rep 6, 22703 (2016).

42. Holtmaat, A. & Svoboda, K. Experience-dependent structural synaptic plasticity in the mammalian brain. Nature Reviews Neuroscience 10, 647–658 (2009).

43. Wu, Y. et al. Contacts between the endoplasmic reticulum and other membranes in neurons. Proceedings of the National Academy of Sciences 114, E4859–E4867 (2017).

44. Schindelin, J., et al. Fiji: an open-source platform for biological-image analysis. Nature methods 9, 676–682 (2012).

45. Taschenberger, H., Leão, R. M., Rowland, K. C., Spirou, G. A. & Von Gersdorff, H. Optimizing synaptic architecture and efficiency for high-frequency transmission. Neuron 36, 1127–1143 (2002).

46. Yang, Y.-M. et al. Septins regulate developmental switching from microdomain to nanodomain coupling of Ca 2+ influx to neurotransmitter release at a central synapse. Neuron 67, 100–115 (2010).

47. Thomas, C. I. et al. Presynaptic Mitochondria Volume and Abundance Increase during Development of a High-Fidelity Synapse. J. Neurosci. 39, 7994–8012 (2019).

48. Lowe, D. G. Distinctive Image Features from Scale-Invariant Keypoints. International Journal of Computer Vision 60, 91–110 (2004).

49. Novikov, A., Major, D., Wimmer, M., Lenis, D. & Bühler, K. Deep Sequential Segmentation of Organs in Volumetric Medical Scans. IEEE Trans. Med. Imaging 38, 1207–1215 (2019).

50. Gray, E. G. Axo-somatic and axo-dendritic synapses of the cerebral cortex. J Anat 93, 420–433 (1959).

51. Diedenhofen, B. & Musch, J. cocor: A Comprehensive Solution for the Statistical Comparison of Correlations. PLOS ONE 10, e0121945 (2015).

52. Wanner, A. A., Kirschmann, M. A. & Genoud, C. Challenges of microtome-based serial block-face scanning electron microscopy in neuroscience. Journal of microscopy 259, 137–142 (2015).

53. Dobrunz, L. E. Release probability is regulated by the size of the readily releasable vesicle pool at excitatory synapses in hippocampus. International Journal of Developmental Neuroscience 20, 225–236 (2002).

54. Vasileva, M., Horstmann, H., Geumann, C., Gitler, D. & Kuner, T. Synapsin-dependent reserve pool of synaptic vesicles supports replenishment of the readily releasable pool under intense synaptic transmission. European Journal of Neuroscience 36, 3005–3020 (2012).

55. Ko, J. et al. Neuroligin-1 performs neurexin-dependent and neurexin-independent functions in synapse validation. The EMBO Journal 28, 3244–3255 (2009).

56. Quinn, D. P. et al. Pan-neurexin perturbation results in compromised synapse stability and a reduction in readily releasable synaptic vesicle pool size. Sci Rep 7, 42920 (2017).

57. Dahlhaus, R. et al. Overexpression of the cell adhesion protein neuroligin-1 induces learning deficits and impairs synaptic plasticity by altering the ratio of excitation to inhibition in the hippocampus. Hippocampus 20, 305–322 (2010).

58. Krueger, S. R., Kolar, A. & Fitzsimonds, R. M. The Presynaptic Release Apparatus Is Functional in the Absence of Dendritic Contact and Highly Mobile within Isolated Axons. Neuron 40, 945–957 (2003).

59. Staras, K. et al. A Vesicle Superpool Spans Multiple Presynaptic Terminals in Hippocampal Neurons. Neuron 66, 37–44 (2010).

60. Alabi, A. A. & Tsien, R. W. Synaptic Vesicle Pools and Dynamics. Cold Spring Harb Perspect Biol 4, a013680 (2012).

61. Denker, A. & Rizzoli, S. Synaptic Vesicle Pools: An Update. Frontiers in Synaptic Neuroscience 2, (2010).

62. Herzog, E. et al. In Vivo Imaging of Intersynaptic Vesicle Exchange Using VGLUT1Venus Knock-In Mice. J. Neurosci. 31, 15544–15559 (2011).

63. Ostroff, L. E., Cain, C. K., Jindal, N., Dar, N. & Ledoux, J. E. Stability of presynaptic vesicle pools and changes in synapse morphology in the amygdala following fear learning in adult rats. Journal of Comparative Neurology 520, 295–314 (2012).

64. Rothman, J. S., Kocsis, L., Herzog, E., Nusser, Z. & Silver, R. A. Physical determinants of vesicle mobility and supply at a central synapse. eLife 5, e15133 (2016).

65. Trotter, J. H. et al. Synaptic neurexin-1 assembles into dynamically regulated active zone nanoclusters. Journal of Cell Biology 218, 2677–2698 (2019).

66. Öztürk, Z., O’Kane, C. J. & Pérez-Moreno, J. J. Axonal Endoplasmic Reticulum Dynamics and Its Roles in Neurodegeneration. Frontiers in Neuroscience 14, (2020).

